# Simultaneous tracking of near-isogenic bacterial strains in synthetic *Arabidopsis* microbiota by chromosomally-integrated barcodes

**DOI:** 10.1101/2023.04.20.537712

**Authors:** Jana Ordon, Julien Thouin, Ryohei Thomas Nakano, Ka-Wai Ma, Pengfan Zhang, Bruno Huettel, Ruben Garrido-Oter, Paul Schulze-Lefert

## Abstract

DNA amplicon-based microbiota profiling currently relies on polymorphisms in microbial marker genes to estimate species diversity and abundance. However, this technique cannot resolve genetic differences among microbial individuals of the same species. We report here the development of modular bacterial tags (MoBacTags) encoding DNA barcodes. These tags facilitate tracking of near-isogenic bacterial commensals in synthetic communities (SynComs), which allow assessment of the contributions of individual bacterial genes to root microbiota establishment in *Arabidopsis thaliana*. Chromosomally-integrated DNA barcodes are co-amplified with endogenous marker genes of the community by integrating corresponding primer binding sites into the barcode. We generated MoBacTag-labeled strains of wild-type *Pseudomonas capeferrum* WCS358 and of pqqF and cyoB mutants with known defects in gluconic acid-mediated host immunosuppression and validated reduced root colonization of both mutants in a 15-member synthetic microbiota. We detected a reduced SynCom load on roots in the presence of the WCS358:pqqF mutant, but not WCS358:cyoB, revealing distinct *pqqF* and *cyoB* activities in a community context. Using MoBacTags, we also show that WCS358 pqqF mutant-specific colonization and community establishment is not *trans*-complemented by wild-type WCS358. Given that gluconic acid production in *P. capeferrum* is indirectly abolished in the pqqF mutant by disruption of pyrroloquinoline quinone (PQQ) biosynthesis, we propose that drastic changes in the root-associated community result from depletion of the cofactor PQQ, which might serve as a common good during root microbiota establishment. Our proof-of-principle experiments illustrate how MoBacTags can be applied to assess scaling of individual bacterial genetic determinants in the plant microbiota.

## Introduction

In natural environments, above- and below-ground organs of flowering plants are inhabited by taxonomically structured yet diverse multi-kingdom microbial communities referred to as plant microbiota. The primary source inocula of the plant microbiota are soil-dwelling bacteria and fungi, which vary among soil types and explain up to 25% of the variation of these plant-associated microbial assemblages (Massoni et al., 2021). When grown in the same soil and environment, the composition of the microbiota also varies between different plant species, indicative of co-diversification of the microbiota with its plant hosts (Fitzpatrick et al., 2018; Wippel et al., 2021). The establishment of systematic microbiota culture collections from plants grown in natural environments, together with gnotobiotic plant systems, has enabled reconstitution experiments with defined (synthetic) microbial communities of reduced complexity. Researchers are now using these communities to test hypotheses regarding microbiota functions under laboratory conditions (Carlstrom et al., 2019; Kremer et al., 2021; Ma et al., 2022; Vorholt et al., 2017). Members of the microbiota form commensal relationships with and provide beneficial services to the plant host (Trivedi et al., 2020), including mobilization of nutrients from soil for root uptake (Castrillo et al., 2017; Harbort et al., 2020), indirect pathogen protection (Berendsen et al., 2012; Carrion et al., 2019; Duran et al., 2018), and abiotic stress tolerance (Fitzpatrick et al., 2018; Santos-Medellin et al., 2021). Establishment of the plant microbiota is dependent on microbe-microbe and microbe-host interactions. Numerous factors influence the composition of these plant-associated microbial assemblages: edaphic factors such as soil pH or soil-derived nutrients (Castrillo et al., 2017; Harbort et al., 2020), microbial competition or cooperation for plant-derived carbohydrates (Berendsen et al., 2018; Hemmerle et al., 2022), specialized plant- or microbe-derived exometabolites (Getzke et al., 2023; Hu et al., 2018; Huang et al., 2019; Voges et al., 2019), including antimicrobials, or pathogen effectors released by co-colonizing phytopathogens (Snelders et al., 2021; Snelders et al., 2020).

Colonization of plants by commensal bacteria typically activates plant defense responses through recognition of conserved microbial epitopes by pattern recognition receptors of the plant innate immune system. At the same time, approximately 40% of cultured commensal strains of the bacterial root microbiota of *Arabidopsis thaliana* have immunosuppressive activity and blunt certain sectors of the plant innate immune system, ultimately leading to commensal-host homeostasis in synthetic microbiota (Ma et al., 2021; Teixeira et al., 2021; Yu et al., 2019). *PqqF* and *cyoB* were recently identified as critical genetic determinants for immunosuppression and root colonization by the plant growth-promoting rhizobacterium (PGPR) *Pseudomonas capeferrum* (*P. capeferrum*) WCS358 on *A. thaliana* roots (Yu et al., 2019). *PqqF* is essential for the biosynthesis of pyrroloquinoline quinone (PQQ; Wei et al., 2016), which serves as redox-sensitive co-factor of several bacterial dehydrogenases and mediates numerous physiological activities, including antioxidant protection (Misra et al., 2012; Oubrie, 2003). *CyoB* encodes the subunit I of the ubiquinol cytochrome *bo3* oxidase (CYO) complex (Stenberg et al., 2007), acts primarily in the energy-providing aerobic respiratory chain, and might also be involved in the global control of pathways for the assimilation of carbon sources (Morales et al., 2006). *PqqF* and *cyoB* commonly function in the production of gluconic acid and its derivative 2-keto gluconic acid, which are proposed to suppress plant immunity by lowering extracellular pH locally (Yu et al., 2019). Suppression of host immune responses by pathogenic microbes invariably leads to increased pathogen proliferation and was recently reported for some commensal bacteria (Ngou et al., 2022; Yu et al., 2019), raising the question of whether other microbiota members also benefit indirectly from compromised host defenses. However, the impact of host immunosuppression by *P. capeferrum* WCS358 on the structure of root-associated microbial communities remains undefined, as does a possible influence exerted by microbe-microbe interactions on *P. capeferrum* immunosuppressive activity.

Quantitative and cultivation-independent analysis of microbial communities using marker genes, such as the *16S rRNA* gene of bacteria or *internal transcribed spacer* (*ITS*) regions of fungi, typically relies on the detection of natural nucleotide polymorphisms in hypervariable regions of these markers, thereby defining distinct microbial taxa. Computational analysis of marker gene-based amplicon DNA sequencing data generated by PCR with marker gene-specific primers has shifted from clustering similar reads into operational taxonomic units (OTUs) to error correction approaches that account for individual amplicon sequence variants (ASVs, Callahan et al., 2016; Peng & Dorman, 2021; Zhang et al., 2021). Despite an increased taxonomic resolution of ASV-classified bacterial communities, *16S rRNA*-based profiling is still unable to capture the true within-phylotype genetic variation of the plant microbiota, as strains with identical marker gene sequences may comprise a bacterial population with other polymorphic loci, as demonstrated for Rhizobiales and *Pseudomonas* lineages (Chiniquy et al., 2021; Garrido-Oter et al., 2018; Karasov et al., 2018). In such cases, only cumulative relative *16S rRNA* abundances originating from multiple strains represented by a phylotype can be retrieved (Figure 1A). This imposes limitations on functional microbiota studies as the aforementioned beneficial bacterial traits vary in a strain-specific manner. Studies on genetic variation within a bacterial phylotype are therefore typically limited to host mono-associations with cultured bacterial or fungal strains, competition experiments with antibiotic markers (Macho et al., 2007), or depend on DNA sequencing of strain-differentiating amplicons restricted to a particular taxon (Ashe et al., 2014). Furthermore, amplification of endogenous marker sequences does not allow differentiation between wild-type and mutant strains in a community context, limiting the application of microbial genetics in gnotobiotic systems.

**Figure 1:**
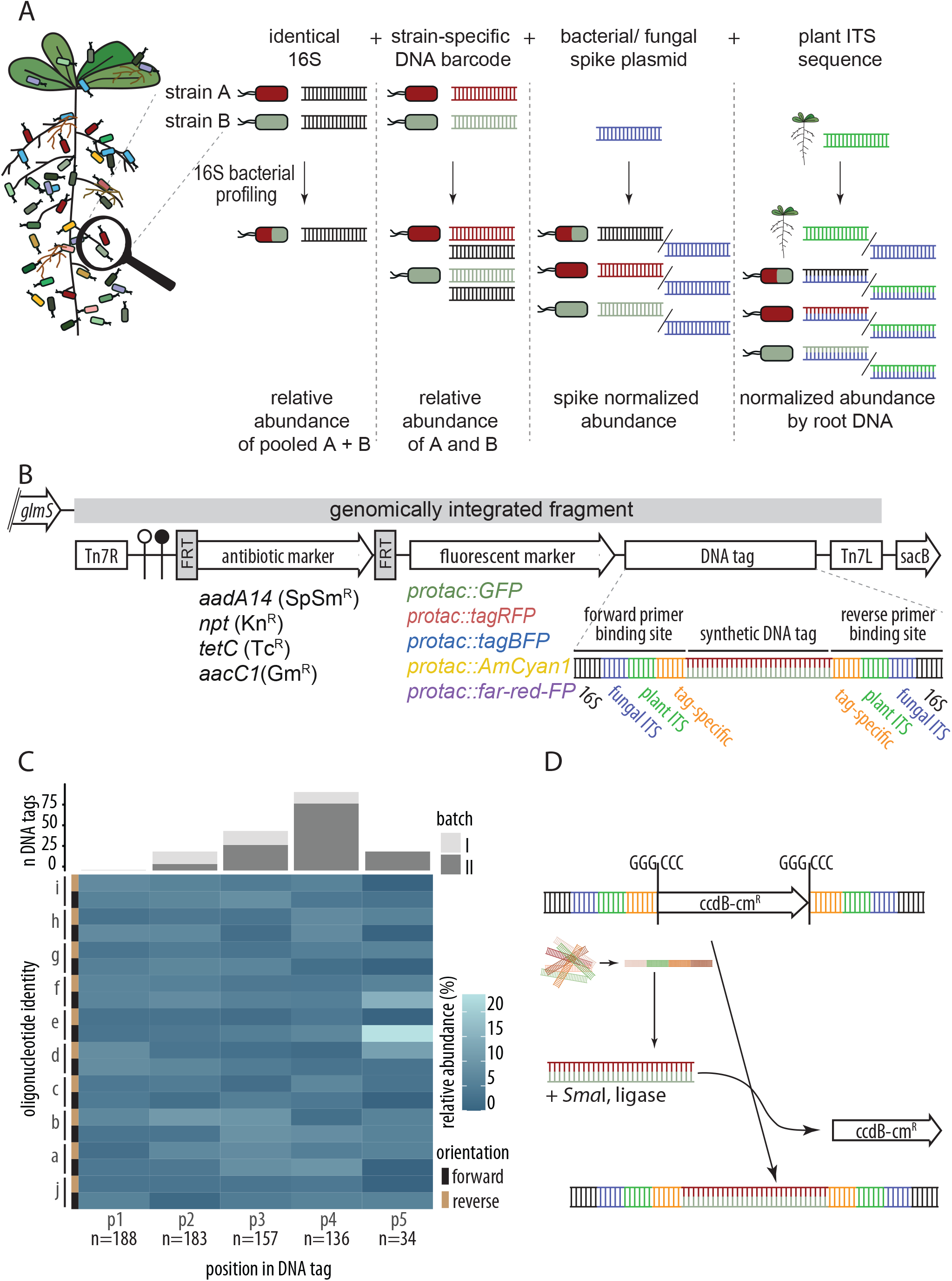
Design and assembly strategy of MoBacTag vectors for simultaneous tracking of near-isogenic bacterial strains. **(A)** Principles of MoBacTag DNA barcodes for discriminating bacterial strains encoding identical *16S rRNA* sequences during amplicon sequencing for microbiota profiling and to quantify microbial load using MoBacTag-based spike-in plasmids. **(B)** Schematic representation of the MoBacTag. Each vector encodes one of the 20 possible combinations of the four antibiotics with the five fluorescent markers. The MoBacTag fragment (grey bar) is chromosomally integrated downstream of the *glucosamine-6-phosphate synthetase* (*glmS*) gene. The synthetic DNA tag is flanked by primer binding sites for amplification of bacterial *16S*, fungal *ITS*, plant *ITS* or barcode sequences. Tn7R, Tn7L: right and left Tn7 borders; FRT: Flp Recombinase Target; SpSmR: Spectinomycin/Streptomycin resistance; KnR: Kanamycin resistance; TcR: Tetracycline resistance; GmR: Gentamycin resistance; *protac*: synthetic bacterial hybrid promoter; xFP: fluorescent protein; *sacB*: *levansucrase* gene conferring sucrose sensitivity. The elements are not drawn to scale. **(C)** Oligonucleotide composition of the synthetic DNA tags. Number of oligonucleotides integrated into the DNA barcode tags (upper panel). Grey colors represent two different assembly batches. Heat map-visualized frequencies of individual oligonucleotides (a-j) in forward or reverse orientation (black, brown) at each position within the DNA barcode tag (lower panel). **(D)** Integration strategy of pre-assembled DNA barcode tags into MoBacTag recipient vectors. Pre-assembled DNA barcode tags are inserted between primer binding sites in exchange for the dominant negative selection marker *ccdB*.

Cellular barcodes can overcome resolution limitations by labeling strains or individual cells with unique DNA sequences, here referred to as DNA barcodes (Figure 1A) (Kebschull & Zador, 2018). DNA barcoding has been used to track cell lineages during experimental evolution (Levy et al., 2015), neuron dispersal (Walsh & Cepko, 1992), stem cell differentiation (Lu et al., 2011), or the development of drug resistance in cancer cells (Bhang et al., 2015). In plant-microbe interactions, DNA barcodes have been used mainly for screening bacterial mutant libraries based on Bar-Seq (Cole et al., 2017; Luneau et al., 2022; Wetmore et al., 2015) and to distinguish closely related *Pseudomonas* strains in the phyllosphere of *A. thaliana* (Shalev et al., 2022). However, simultaneous profiling of taxonomically diverse plant-associated microbial communities by amplicon sequencing of marker genes and of DNA barcodes of near-isogenic strains is currently not possible.

Here, we describe a **Mo**dular **Bac**terial **Tag** (MoBacTag) tool to label a broad range of taxonomically distinct bacteria with a DNA barcode as well as a fluorescent tag. The DNA barcode allows discrimination of bacteria that cannot be distinguished when profiling microbial communities using conventional marker gene amplicon sequencing. Since the DNA barcode is flanked by *V5*–*V7 16S rRNA*-specific, fungal *ITS*-specific, plant *ITS*-specific, and barcode-specific primer binding sites, the abundances of barcoded bacteria can be determined simultaneously with abundances of unlabeled bacteria and fungi by amplicon sequencing of the corresponding marker genes. Furthermore, DNA barcode-harboring plasmids are used as spike DNA to estimate microbial load and calculate the ratio of plants to microbes. As a proof of principle, we recapitulated the colonization defect of *P. capeferrum* WCS358 *cyoB* and *pqqF* immunosuppressive mutants with MoBacTags. By simultaneously analyzing DNA barcodes and *16S rRNA* sequences, we reveal an activity specific to the WCS358 *pqqF* mutant in community establishment, leading to a reduction of total bacterial load on *A. thaliana* roots. As wild-type WCS358 does not *trans*-complement the reduced SynCom load in presence of the WCS358 pqqF mutant on roots, we propose that the cofactor PQQ, which might be depleted by the WCS358:pqqF mutant, serves as a common good during root microbiota establishment.

## Results

### Design of MoBacTag tools

To track bacteria with identical marker gene sequences during community profiling, we developed MoBacTag tools encoding a unique artificial sequence identifier, termed strain-specific barcode DNA tag, transformed into bacterial strains of interest. To overcome potential positional bias resulting from random chromosomal integration, MoBacTag tools have been designed for orientation-specific high-frequency insertion into bacterial chromosomes at the *Tn7* attachment sites (Peters & Craig, 2001). These sites are located in the neutral, intergenic region downstream of the *glmS* gene encoding the essential *glucosamine-6-phosphate synthetase* (Craig, 1996). We identified *glmS* orthologues in 98.6% (426/432) of all bacterial draft genomes in the *Arabidopsis*-derived bacterial culture collection (*At*-SPHERE; Bai et al., 2015) and 90% (384/426) encode a single copy of *glmS* and thus contain a unique *Tn7* attachment site. The site-specific chromosomal integration of *Tn7*-mediated by *tnsD* depends on the recognition site encoded in the last 36 nts of the *glmS* open reading frame. The alignment of *At*-SPHERE commensal-encoded *tnsD* recognition sites revealed that the first two positions of each codon, which are important for binding *tnsD*, are highly conserved in the *At*-SPHERE culture collection (Supplemental figure 1; Mitra et al., 2010). Thus, the MoBacTag tools are *per se* applicable to strains representing the majority of taxa that define the core bacterial microbiota of plants.

The chromosomally integrated fragment contains (*i*) the minimal *Tn7* elements (Tn7R, Tn7L) (Choi & Schweizer, 2006), (*ii*) two terminator sequences, (*iii*) an antibiotic marker flanked by yeast *Flp* recombinase recognition sites for sequential excision, (*iv*) a fluorescent marker and (*v*) a barcode DNA tag (Figure 1B). Regulatory elements, expression cassettes, and the barcode DNA tag were first individually mobilized into level 1 vectors using modular cloning principles (Supplemental table 3; Weber et al., 2011). All 20 possible combinations of the four antibiotics and five fluorescent markers were assembled into a modular cloning-adapted pSEVA211-based backbone (Silva-Rocha et al., 2013). For each antibiotic–fluorescent marker combination, we constructed three to five ready-to-use vectors with distinguishable barcode DNA tags (Supplemental table 5). Chromosomal integration into *attTn7* is enforced by the pSEVA211 vector backbone containing the restrictive *R6K* origin of replication (Miller & Mekalanos, 1988), which renders the vector unstable in most bacteria, and the negative selection marker *sacB* (Supplemental figure 2).

For profiling of microbial communities by amplicon sequencing, the barcode DNA tag is flanked by conserved bacterial *V5*–*V7 16S rRNA* sequences, conserved fungal *ITS1/ ITS2* sequences, plant *ITS p4/p5* sequences and barcode-specific primer binding sites (Figure 1B). MoBacTag plasmid DNA was additionally used as spike-in DNA during library preparation for amplicon sequencing and comparative analysis. As the *16S*, fungal *ITS*, and plant *ITS* primer binding sites flank the barcode DNA tag, read counts assigned to chromosomally integrated barcode DNA tags, bacterial *16S rRNA* or fungal *ITS* can be normalized to spike-in barcode DNA tag-specific read counts. These ratios can then be used to calculate bacterial to plant, fungal to plant or bacterial to fungal ratios. Finally, these ratios provide an estimate of the microbial load of the corresponding microbial kingdom in the sampled plant compartment (Figure 1A).

Unique barcode DNA tags were generated by random blunt-end ligation of an equimolar ratio of ten different, double-stranded oligonucleotides (38 nts), each consisting of four pyrosequencing-friendly barcodes, followed by DNA fragment size selection. The final barcode DNA tags preferentially consisted of an array of four oligonucleotides, resulting in an average length of approximately 150 nts per barcode (Figure 1C). As the position and orientation of each oligonucleotide within the array appeared to be random, this method could theoretically generate 160,000 different barcode DNA tags for an array length of four oligonucleotides (Figure 1C). By incorporating additional barcode oligonucleotides into the ligation reaction, the total number of unique barcode DNA tags can be further increased. The pre-assembled barcode DNA tags were then integrated by restriction/ligation (using *Sma*I) in exchange for a negative *ccdB* selection cassette previously integrated between primer binding sites of recipient vectors (Figure 1D). As *Sma*I recognition sites are only disrupted by the integration of a barcode DNA tag, plasmids with an integrated barcode DNA tag were enriched over multiple restriction/ligation cycles.

### Validation of barcode DNA tags as spike and artificial *16S rRNA* sequence

Correlation of read counts with the spike DNA concentration was tested for each oligonucleotide pair in root and peat matrix samples to ensure that spike DNA abundance was reflected in barcode DNA tag-specific read counts (Figure 2A–D). Spike-specific barcode read counts from amplicon sequencing with *16S rRNA* (A), fungal *ITS* (B), plant *ITS* (C), or tag-specific (D) primers correlated linearly over five orders of magnitude with spike plasmid DNA concentrations in both root and peat matrix samples. This suggests that spike-specific read counts can be used to normalize *16S rRNA*, fungal *ITS*, plant *ITS,* and chromosomally integrated barcode DNA tag read counts.

**Figure 2:**
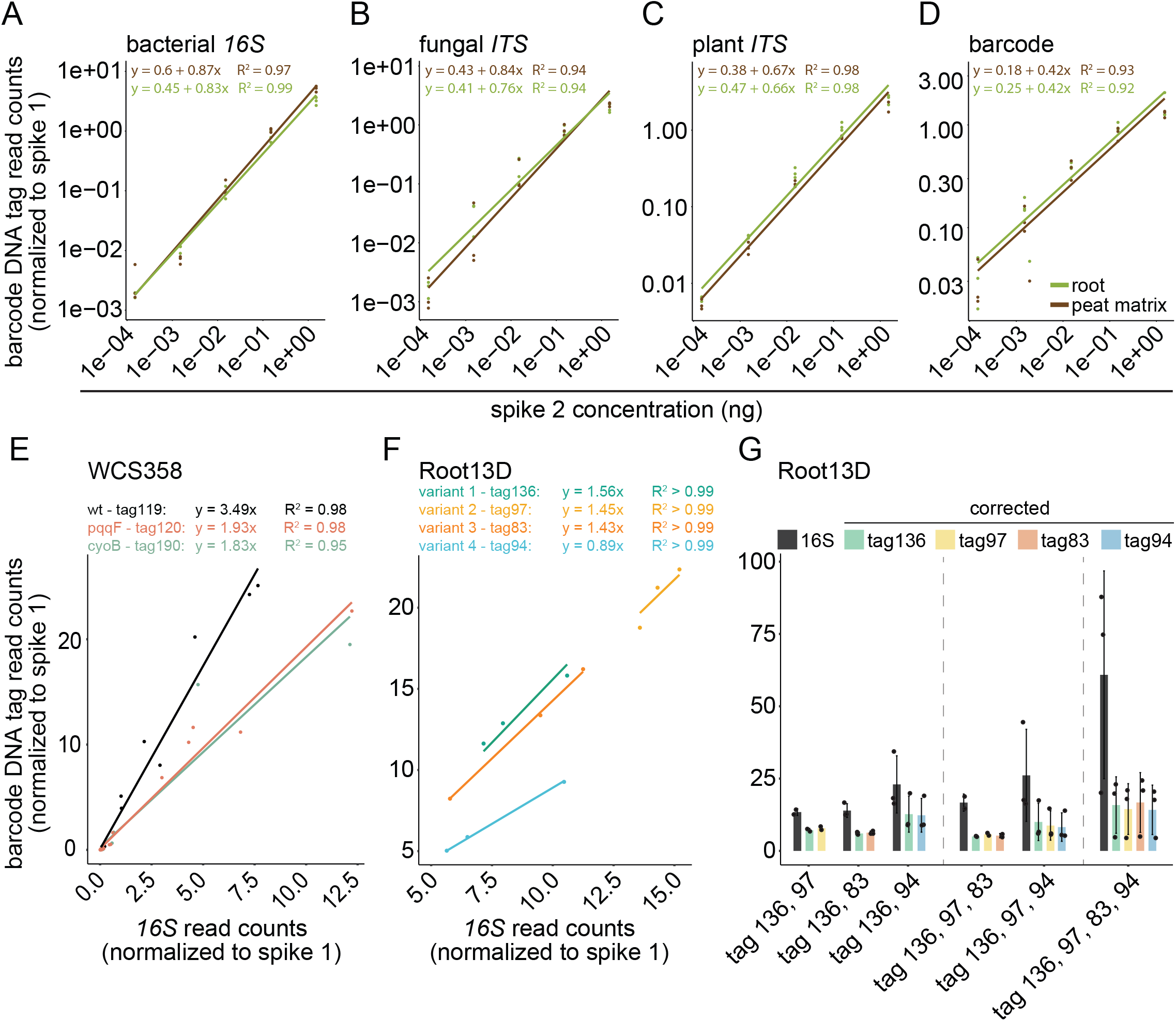
Validation of the MoBacTag for spike normalization and strain abundance estimates during amplicon sequencing. **(A–D)** Linear correlation of normalized spike-specific read counts with spike concentrations obtained from amplicon sequencing of root (n=15) and peat (n=15) matrix samples using bacterial *16S*- (799/1192; A), fungal *ITS*- (ITS1/ ITS2; B), plant *ITS*- (ITS-p4/ ITS-p5; C) and barcode- (JT259/ JT262; D) specific primers. **(E)** Linear correlation of normalized barcode-specific read counts with normalized *16S*-specific read counts for correction factor adjustment of individual chromosomally integrated barcodes into WCS358 (n=16 per variant, corresponding to n=10 from root, n=3 from peat matrix and n=3 from input). **(F)** Linear correlation of normalized barcode-specific read counts with normalized *16S*-specific read counts for correction factor adjustment of individual chromosomally integrated barcodes into Rhizobiales Root13D (n=3 per variant). **(G)** Correction factor-adjusted and normalized frequencies of barcode and 16S rRNA-specific read counts within samples with increasing numbers of equimolar mixed Rhizobiales Root13D barcode strains. The colors indicate the DNA barcode tags.

A barcode DNA tag varies in length and structure not only from natural endogenous *V5*–*V7 16S rRNA* gene sequences, but also slightly between different barcode DNA tags. We compared the efficiencies of PCR amplification and DNA sequencing for natural *16S rRNA* with corresponding barcode DNA tags. To exclude possible biases due to known variations in *16S rRNA* copy numbers in different bacterial taxa (Vetrovsky & Baldrian, 2013), different MoBacTags were inserted at identical *Tn7* chromosomal integration sites into the same *16S rRNA* genetic background of *Pseudomonas capeferrum* WCS358 (termed tag119, tag120 and tag190). Short-read sequencing of individual MoBacTag-labeled bacteria revealed 3.5-fold higher read counts for tag119, 1.9-fold higher for tag120 and 1.8-fold higher for tag190 compared to the corresponding natural endogenous *16S rRNA* read counts of *P. capeferrum* WCS358 (Figure 2E). These data indicate that despite identical chromosomal integration site, primer binding sites and *16S rRNA* genetic background, individual barcode DNA tags exhibit different tag-to-*16S* read count ratios. We also generated and tested four variants of the root commensal Rhizobiales Root13D with MoBacTags integrated at different loci through homologous recombination. The data also showed tag-specific amplification with read counts that were 1.6-fold higher for tag104, 1.5-fold higher for tag93 and 1.4-fold higher for tag94 compared to the corresponding natural endogenous *16S rRNA* read counts (termed tag104, tag93, tag94; Figure 2F). Fewer read counts were obtained for DNA tag91 compared to endogenous *16S rRNA* (Fig. 2F), indicating tag-specific amplification bias for MoBacTags at identical and different chromosomal integration sites. To ensure that the read counts obtained from different barcode DNA tags accurately reflect the respective bacterial abundance, the tag-to-*16S* count ratios were used as a correction factor to the barcode DNA tag-specific read counts. By sequencing of individual bacterial cultures as exemplified by Rhizobiales Root13D or communities, as shown for WCS358, in which the barcode reads are correlated with the endogenous *16S rRNA* reads, barcode tag-specific correction factors for MoBacTag-labelled strains can be estimated. We then investigated potential combinatorial effects of multiple MoBacTags within a sample. Genomic DNA of Root13D strains carrying the four different barcode DNA tags were mixed in a 1:1 ratio with increasing complexity, i.e., two, three or all four strains (Figure 2G). Similar read counts were obtained for each tag per condition when using the tag-specific correction factors determined from the ratios of tag to *16S* reads. In summary, the number of MoBacTag-labeled strains within a condition does not change the number of tag-specific reads per strain, so the corrected tag-specific reads reflect strain abundance.

### MoBacTag-labeling of taxonomically different root microbiota members

As with any *Tn7*-mediated insertion, genome integration of the MoBacTag into the bacterial strain of interest requires four (not necessarily consecutive) working days, assuming that a DNA transformation protocol has already been established for the bacterium of interest (Figure 3A) (Choi & Schweizer, 2006). For conjugation-based tagging of root-derived *Rhodanobacter* spp. Root179, *Tn7* attachment sites and natural antibiotic resistances were investigated. For conjugation, MoBacTag multigene vectors were mobilized into auxotrophic, conjugation-competent *E. coli* BW29427 (Dehio & Meyer, 1997). In addition to selecting transformants based on MoBacTag-encoded antibiotic resistance, selection against antibiotic-resistant *E. coli* BW29427 was achieved by depletion of diaminopimelic acid (DAP), which is required for the survival of *E. coli* BW29427 but not the target strains (Wang et al., 2015). The presence of the MoBacTag fragment was verified by PCR of bacterial colonies using plant *ITS*-specific oligonucleotides (Figure 3B, JT103/JT108). The absence of PCR-amplified fragments from non-transformed bacterial colonies confirmed the specific binding of the plant *ITS* oligonucleotides to the MoBacTag in *Rhodanobacter* R179. In community profiling experiments with 15 bacterial strains selected from the *At*-R-SPHERE collection (SynCom modified from Wippel et al., 2021), no bacteria-derived sequences were recovered using the plant *ITS*-specific oligonucleotides, supporting the target specificity of the plant *ITS* primers. In addition, transformants were genotyped by PCR using *16S rRNA*-specific oligonucleotides, resulting in two size-separable PCR products, as the barcode DNA tag is shorter than the endogenous *16S rRNA* amplicon (Figure 3B, 799/1192). Chromosomal integration was validated using a combination of *Tn7*-specific and *glmS* strain-specific oligonucleotides (Figure 3B; Choi & Schweizer, 2006). Further PacBio long-read genome re-sequencing of MoBacTag-labeled and wild-type *Arabidopsis* root-derived Rhizobiales strains revealed no chromosomal rearrangement induced by the *Tn7*-mediated insertion (Supplemental figure 3A). Thus, the tagged strains differ from the wild type only by the chromosomally integrated MoBacTag, which could be validated by PCR-based genotyping. Comparisons of unlabeled and MoBacTag-labeled strains showed indistinguishable colony morphologies and growth curves in liquid medium (Supplemental figure 2B).

**Figure 3:**
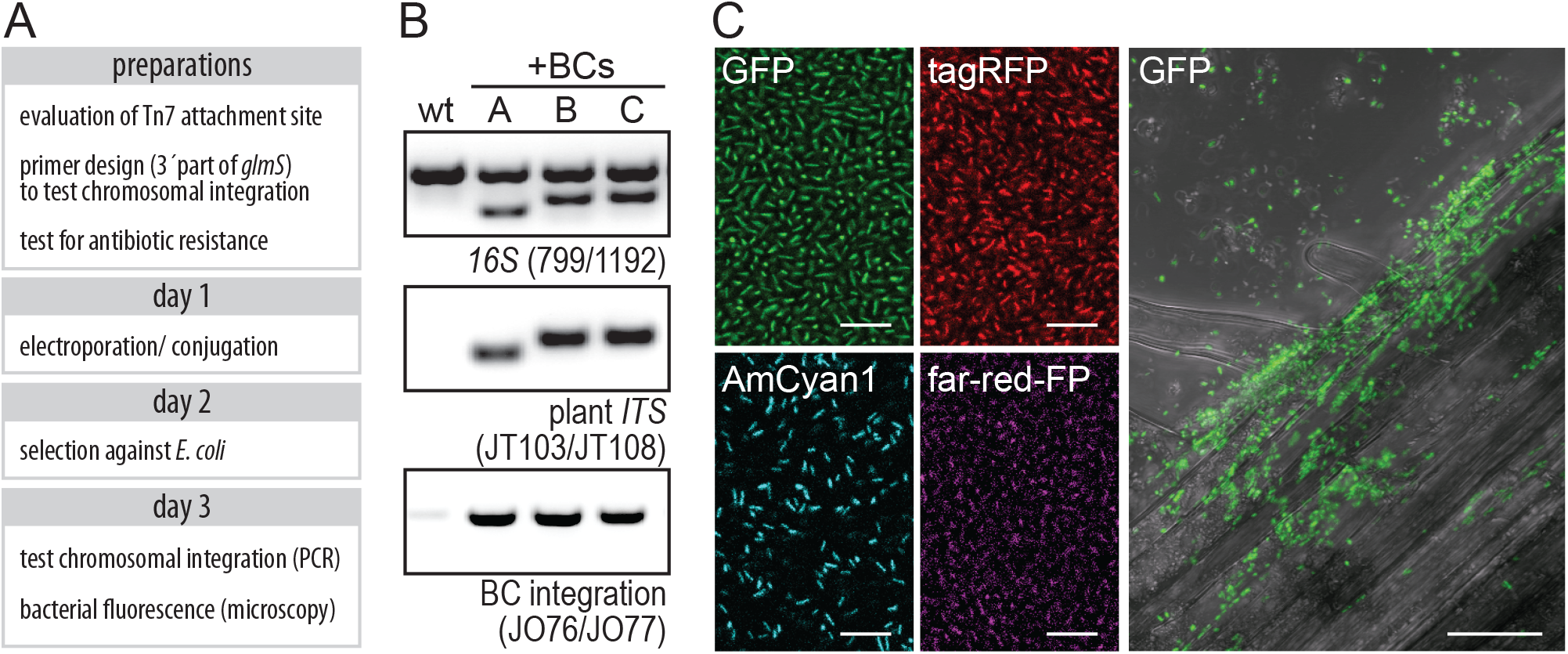
Chromosomal integration of MoBacTags and expression of fluorescent tags. **(A)** General timeline to label a target strain by chromosomal MoBacTag integration. Depending on the target strain, *Tn7*-mediated insertion requires four (not necessarily consecutive) working days. **(B)** Validation of MoBacTag transformation and chromosomal integration. DNA was extracted from transformed (+BC) and un-transformed *Rhodanobacter* Root179 strains, PCRs were performed using indicated primers and amplicons were visualized on an agarose gel. To test barcode integration, *Tn7* element- and *glmS*-specific primers were used. **(C)** MoBacTag-labeled strains express fluorescent proteins. Expression of MoBacTag-encoded fluorescent protein was detected using live confocal laser scanning microscopy in liquid medium and on *A. thaliana* roots. Scale of bacteria in liquid culture: 10 µm, scale on roots: 20 µm.

MoBacTags encoding different fluorescent markers were transformed into the root-derived commensal *Rhodanobacter* R179. Live cell imaging by confocal laser scanning microscopy was used to detect the expression of the four chromosomally-encoded fluorescent markers driven by the *tac* promoter in bacteria cultivated on nutrient-rich medium or in association with the plant host (Figure 3C). This observation corroborates early studies on the *tac* promoter that showed that it can drive efficient gene expression in several families of Proteobacteria (Deboer et al., 1983; Morales et al., 1991). Differences in fluorescence intensities were observed for different tagged strains, when grown in nutrient-rich medium or on roots and with the different fluorophores. In particular, the GFP and AmCyan1 proteins yielded stronger fluorescence signals. Expression of fluorescent proteins was also detected in MoBacTag-labeled *Pseudomonas capeferrum* WCS358, *Pseudomonas simiae* WCS417, *Xanthomonas campestris* pv. *vesicatoria*, and *Rhizobiales* spp. Root13D, indicating a robust activity of the *tac* promoter in the corresponding bacterial taxa (Supplemental figure 3C).

### Response of a root-associated community to a MoBacTag-labeled commensal

To investigate whether a chromosomally integrated MoBacTag alters bacterial community structure in root or peat matrix compartments, microbiota reconstitution experiments were performed using gnotobiotic *Arabidopsis* seedlings grown in Jiffy-based gnotopots with defined (synthetic) bacterial communities (SynCom; Kremer et al., 2021). A taxonomically diverse SynCom consisting of 15 *Arabidopsis* root-derived bacteria from the *At*-R-SPHERE culture collection was co-inoculated with *Pseudomonas capeferrum* WCS358 wild-type or the MoBacTag-labeled variant WCS358:BC (SynCom modified from Wippel et al., 2021). As strain Root418 was absent or detectable only at low abundances in output communities (<4%), we depleted Root418 sequence reads *in silico* prior to community analyses. As all SynCom members were distinguishable on the basis of their endogenous *V5*–*V7 16S rRNA* sequences, Principal Coordinate Analyses (PCoA) of Bray–Curtis dissimilarities of *V5*–*V7 16S rRNA* reads normalized to spike reads were performed to assess potential differences in bacterial community composition in the presence of *P. capeferrum* WCS358 wild type or its MoBacTag-labeled variant. Most of the variation was explained by compartments, separating root and peat matrix-associated communities (64% of variation in the first component; Figure 4A–C), which suggests that live plant roots exert a major influence on SynCom composition. We expected significant differences in root- and peat matrix-derived bacterial communities in the presence of wild-type *P. capeferrum* WCS358 or the barcoded variant WCS358:BC since the MoBacTag is treated as an additional strain/*V5*–*V7 16S rRNA* sequence (Figure 4A). Indeed, the differences in community composition decreased after *in silico* depletion of MoBacTag- and WCS358 *16S rRNA*-specific reads (Figure 4B,C). Thus, the bacterial community harboring a MoBacTag-labeled WCS358 strain is very similar to the community containing the untagged WCS358 wild-type strain.

**Figure 4:**
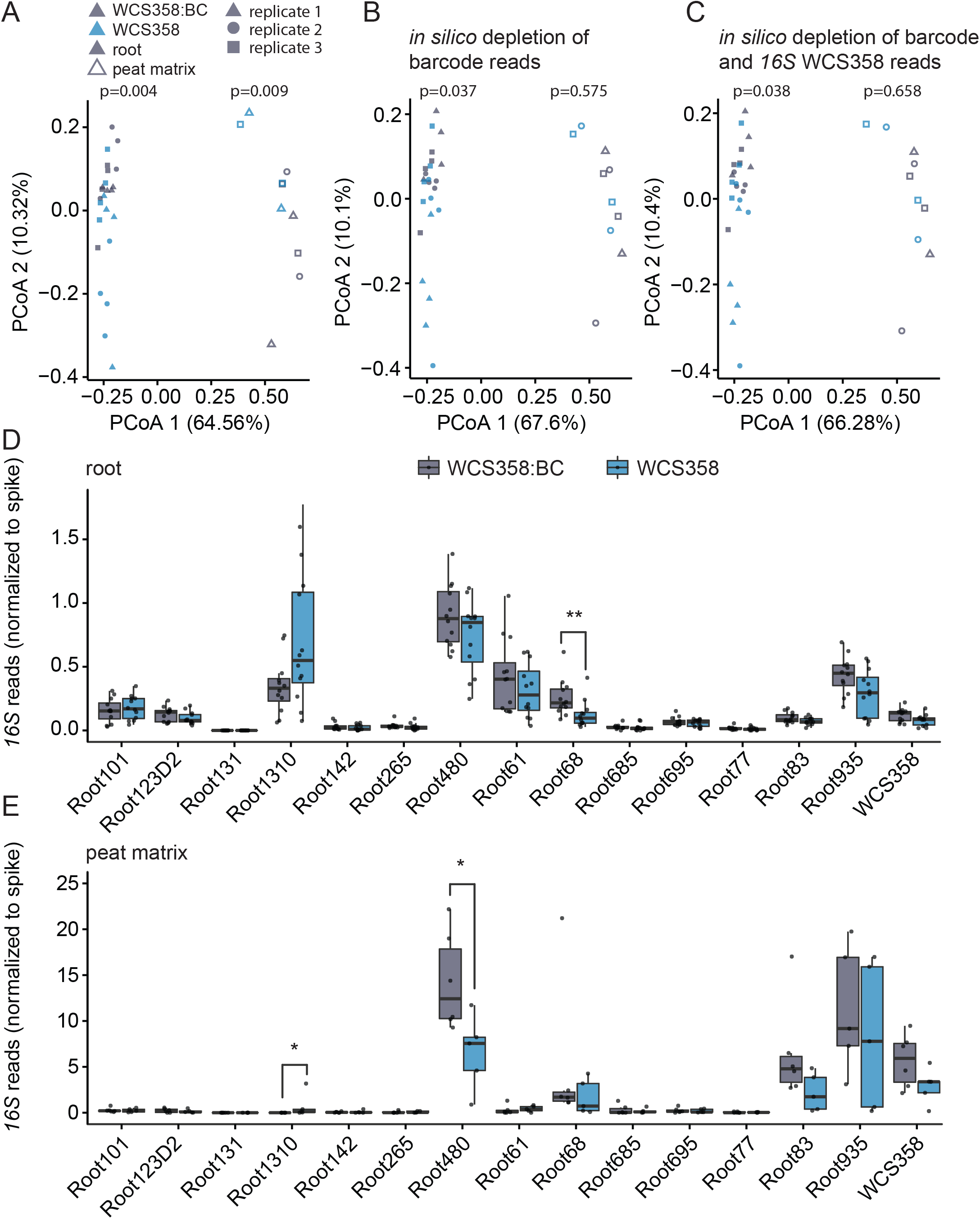
The influence of MoBacTag on root microbiota establishment. **(A–C)** Coordination of the spike-normalized reads from a 15-member synthetic community with wild-type or MoBacTag-labeled *P. capeferrum* WCS358 in the *A. thaliana* root (n=24) and peat matrix (n=10) compartment (A), upon *in silico* depletion of barcode reads (B) and additional *in silico* depletion of WCS358 *16S rRNA* reads (C). Shapes represent the compartment and colors represent WCS358 derivative. n values indicate biological samples collected from three independent replicates. Ellipses correspond to Gaussian distributions fitted to each cluster (95% confidence interval). P values indicate statistical significance determined using a permutational analysis of variance (PERMANOVA) test between communities including the labeled or unlabeled WCS358 derivate (permutation = 999, P < 0.05). **(D,E)** The normalized abundance of individual strains in the root (n=24) (D) and peat matrix (n=10) (E) compartments upon co-inoculation with the labeled (WCS358:BC) or unlabeled (WCS358) strain. Asterisks indicate statistical significance determined using Dunn’s test: * = p<0.05; ** = <0.01. The box plots center on the median and extend to the 25th and 75th percentiles, and the whiskers extend to the furthest point within 1.5x the interquartile range.

Consistent with the Bray–Curtis dissimilarity analyses, the abundance of individual bacterial community members in root and peat matrix compartments were very similar in communities formed in the presence of untagged WCS358 or the MoBacTag-labeled variant WCS358:BC (Figure 4D and 4E). A few SynCom members diverged in abundance depending on the presence of unlabeled WCS358 or labeled WCS358:BC, such as *Pseudomonas* spp. Root68 in the root compartment (Figure 4D) or *Streptomyces* spp. Root1310 and Xanthomonadales Root480 in the peat matrix compartment (Figure 4E), but these differences were not consistent across independent replicates (Supplemental figure 4). Accordingly, the observed differences are likely due to intrinsic stochastic effects during the establishment of bacterial communities as supported by the analysis of individual, independent experiments (Supplemental figure 3) and do not result from the gentamicin resistance cassette introduced together with the MoBacTag fragment (Figure 1B). In conclusion, the chromosomally integrated MoBacTag fragment in WCS358 had little influence on the establishment of 15-member bacterial communities in both the root and peat matrix compartments.

Spike normalization to determine microbial load and community composition *in planta* To investigate whether the immunosuppressive activity of *P. capeferrum* WCS358 selectively promotes its own colonization or also influences root colonization by other members of the bacterial community, wild-type WCS358, WCS358:pqqF or WCS458:cyoB mutants (Yu et al., 2019) were each, or in combination, co-inoculated with the 15 members of the bacterial SynCom described above on germ-free *Arabidopsis* seedlings. For this purpose, wild-type WCS358, WCS358:pqqF and WCS358:cyoB were labeled with MoBacTags differing in DNA barcodes, but an identical antibiotic resistance cassette, so that potential marginal effects of the MoBacTag on community establishment were identical for all conditions (Figure 4).

We first focused on the SynCom members by *in silico* depletion of WCS358 *16S rRNA* reads. Read counts relative to sample read depth appeared to indicate enhanced root colonization of *Streptomyces* spp. Root1310 and decreased colonization of *Microbacterium* spp. Root61 in the presence of WCS358:pqqF mutant compared to the WCS358 wild type (Figure 5A). However, normalization of *16S rRNA* reads to spike or plant *ITS*-derived reads revealed a substantially reduced total bacterial load on roots colonized by the WCS358:pqqF mutant (Figure 5B, 5C). Therefore, the altered relative abundances of *Streptomyces* spp. Root1310 and *Microbacterium* spp. Root61 in the presence of WCS358:pqqF are likely an artefact due to the compositional nature of community profiling in the absence of normalization. As a result, further analyses were only performed on the spike-normalized data.

**Figure 5:**
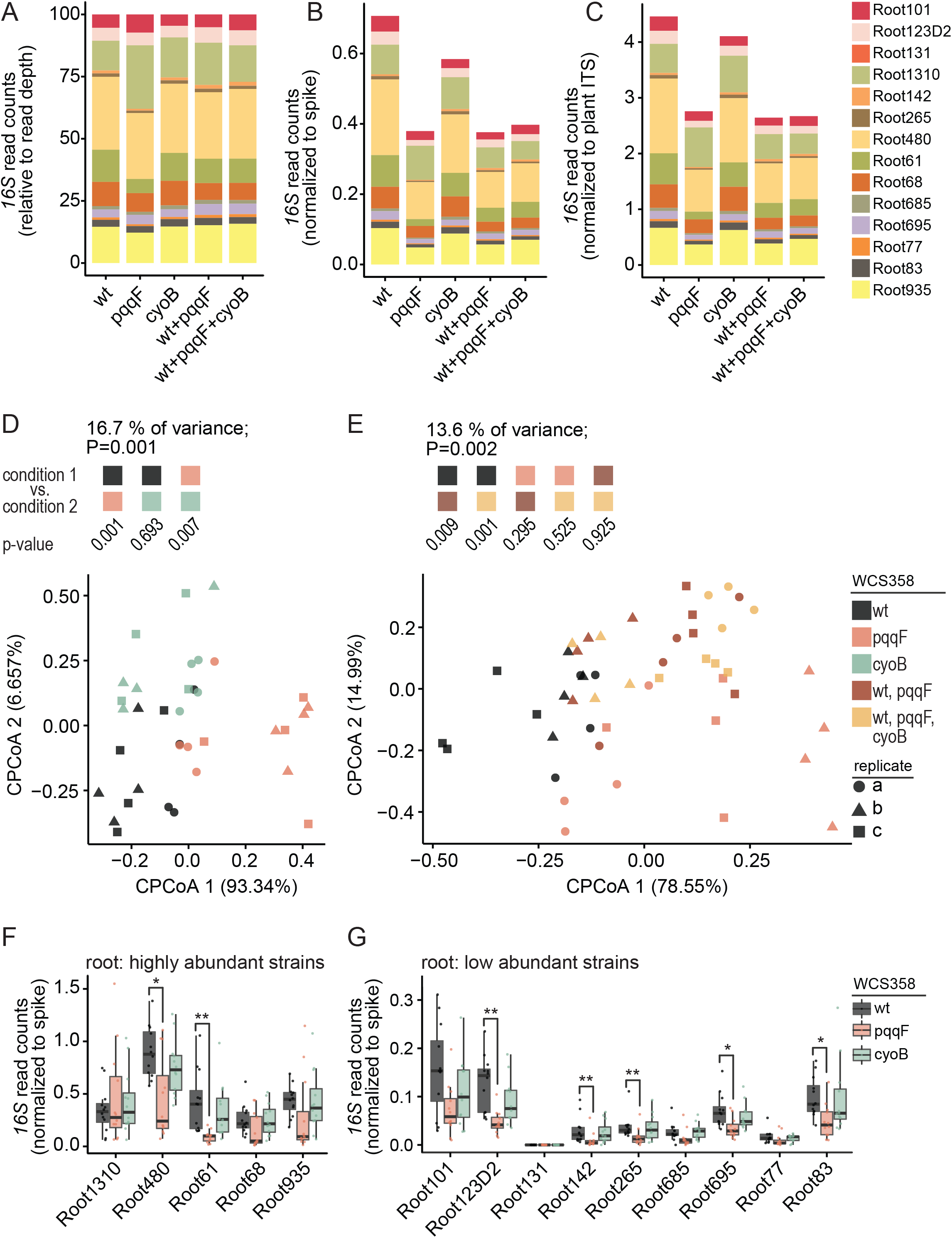
*P. capeferrum* WCS358-encoded pqqF modulates the root microbiota. **(A–C)** Relative (A), spike-normalized (B) and plant ITS-normalized (C) abundance of synthetic community members co-inoculated with *P. capeferrum* WCS358 wild-type, WCS358:pqqF or WCS358:cyoB mutant on *A. thaliana* roots plotted as stacked bar plots. Colors indicate SynCom members. The mean of 12 biological samples collected from three independent replicates is displayed. **(D,E)** Constrained coordination of the microbial profile of a 15-member synthetic community in presence of *P. capeferrum* WCS358 wild type (black, n=12), WCS358:pqqF (red, n=12) or WCS358:cyoB (green, n=12) mutant or WCS358 wild type plus WCS358:pqqF (dark red, n=12) or additional WCS358:cyoB (yellow, n=12) on *A. thaliana* roots. P values indicate statistical significance determined by permutational analysis of variance (PERMANOVA) test between communities co-established with WCS358 derivatives indicated by the colored box (permutation = 999). Shapes represent independent experiments. **(F,G)** The normalized abundance of strains in the root compartment upon co-inoculation with *P. capeferrum* WCS358 wild type (black, n=12), the pqqF (red, n=12) or cyoB (green, n=12) mutant. Asterisks indicate statistical significance determined using Dunn’s test: * = p<0.05; ** = <0.01. The box plots center on the median and extend to the 25th and 75th percentiles, and the whiskers extend to the furthest point within 1.5x the interquartile range.

PCoA of Bray–Curtis dissimilarities of spike-normalized *16S rRNA* reads showed that the reduced SynCom load on roots in the presence of the WCS358:pqqF mutant compared to wild-type WCS358 correlates with the formation of distinct communities (Figure 5D,E). Seven SynCom members showed a significantly reduced root colonization capacity in the presence of WCS358:pqqF compared to WCS358 wild type (Figure 5F,G). The reduced total microbial load can therefore be explained by a similar trend for half of all SynCom members, suggesting that PQQ biosynthesis by wild-type WCS358 presumably indirectly promotes root colonization by taxonomically diverse members of the root microbiota. Unexpectedly, however, root-associated communities established in the presence of wild-type WCS358 plus WCS358:pqqF showed only a slight shift towards communities containing wild-type WCS358 alone (Figure 5B, 5C, 5E). Thus, the presence of the PQQ-deficient WCS358 strain has an unexpected dominant influence on root microbiota establishment that is not complemented by co-inoculation with the PQQ-producing wild-type WCS358. This suggests that immunosuppression mediated by gluconic acid synthesis in wild-type WCS358 is insufficient to support the establishment of wild type-like root communities in the presence of the WCS358:pqqF mutant.

Unlike WCS358:pqqF, the loss of *cyoB* in WCS358 had only a minor effect on the total bacterial load and the composition of the root-associated communities (Figure 5D,F,G). As genetic depletion of *cyoB* in *P. capeferrum* WCS358 eliminates 2-keto-D-gluconic acid production but preserves residual gluconic acid biosynthesis *in vitro* (Yu et al., 2019), the wild-type-like community found here in the presence of WCS358:cyoB might be explained by residual gluconic acid production *in planta*.

### Tracking of near-isogenic commensal strains in microbial community contexts

Next, we asked whether the reduced root colonization by the WCS358:cyoB and WCS358:ppqF mutants reported from previous inoculation experiments in unsterilized soil-sand matrix (Yu et al., 2019), can be reproduced in a peat matrix-based gnotobiotic plant system (Kremer et al., 2021) in the presence of a defined 15-member bacterial community. Consistent with the relative *16S rRNA* read counts for the 15 members of the SynCom (Figure 5A), the interpretation of the *P. capeferrum* WCS358-specific relative *16S rRNA* read counts was limited by the differential total microbial load between conditions (Figure 6A). Accordingly, no difference in abundance was detected between the wild-type WCS358 and the WCS358:pqqF mutant when analyzing the relative read counts. However, the spike-normalized abundances of *16S rRNA* reads from *P. capeferrum* WCS358 strains derived from either wild type, WCS358:cyoB, or WCS358:pqqF mutants recapitulated the reduced root colonization by the mutants in our peat-based gnotobiotic plant system previously observed in an unsterilized soil-sand matrix (Figure 6B). In agreement with published data, the abundance of wild-type WCS358 and WCS358 mutants in the peat matrix compartment was comparable (Figure 6A,B; Yu et al., 2019). Thus, *pqqF* and *cyoB* are needed to specifically promote WCS358 colonization of the root compartment but are dispensable for bacterial growth in the peat matrix.

**Figure 6:**
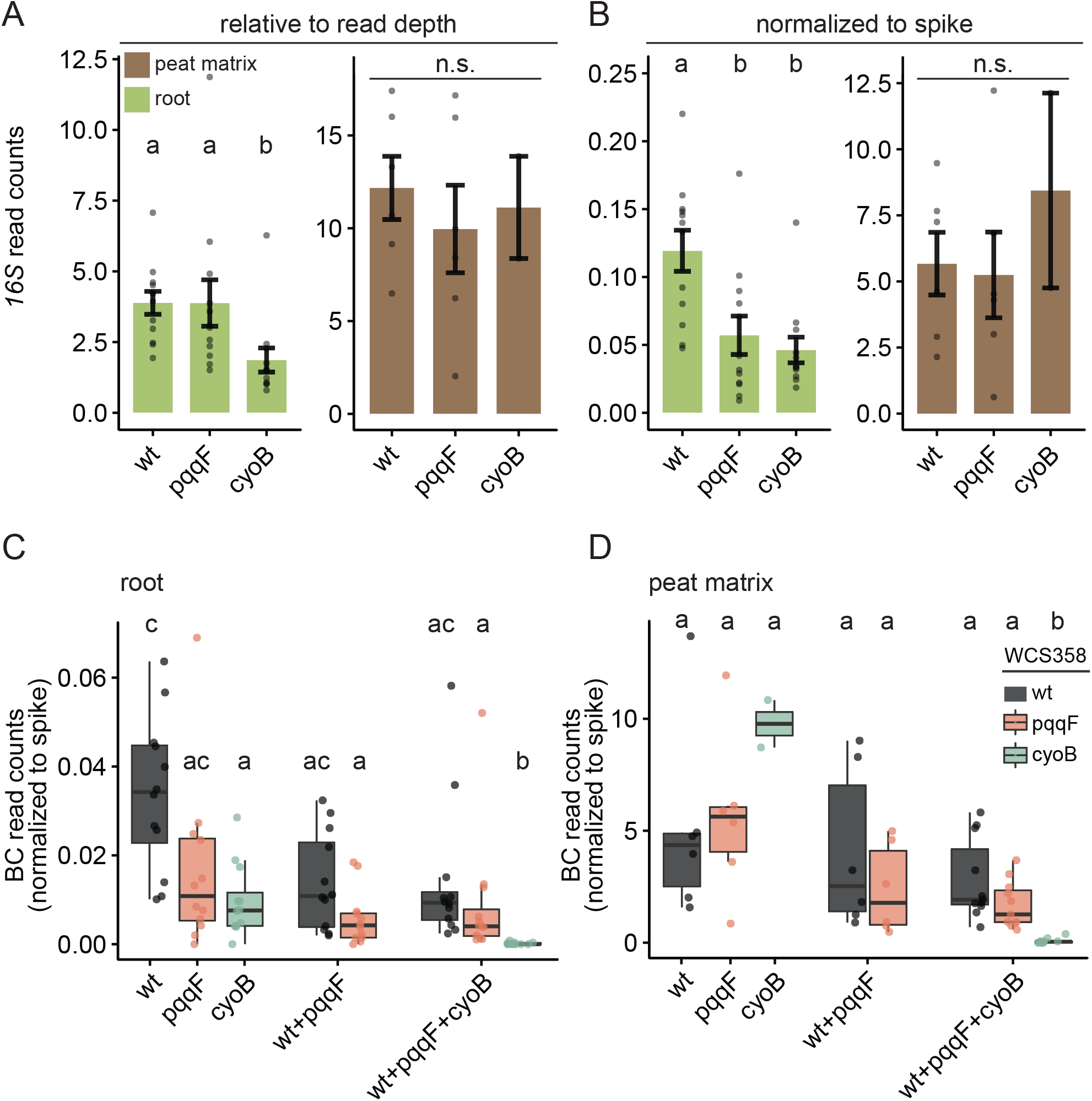
MoBacTags recapitulate the root colonization deficit of WCS358 pqqF and cyoB mutants. **(A,B)** Relative (A) and spike-normalized (B) abundance of *P. capeferrum* WCS358 wild-type, WCS358:cyoB and WCS358:pqqF in the root (green, n=36) and peat matrix (brown, n=14) compartment quantified by *16S rRNA* read-counts. **(C,D)** The normalized abundance calculated using barcode-specific reads of *P. capeferrum* WCS358 wild type (black), the pqqF (red) or cyoB (green) strains in the root (n=60) (C) and peat matrix (n=31) (D) compartment upon co-inoculation with the 15-member synthetic community alone, or in indicated mixtures. Letters indicate statistical significance (p<0.05) determined using Dunn’s test. The box plots center on the median and extend to the 25th and 75th percentiles, and the whiskers extend to the furthest point within 1.5x the interquartile range.

Finally, to test whether strain-resolved abundances of MoBacTags can be retrieved during community profiling of bacteria with identical *16S rRNA* sequences, the abundances of the MoBacTags incorporated into *P. capeferrum* WCS358 wild type, the WCS358:pqqF, and WCS358:cyoB mutants were examined. Spike-normalized, corrected barcode DNA tag reads independently confirmed similarly reduced root and wild type-like peat matrix colonization by the tested WCS358 mutants when only one WCS358 genotype was added to the 15-member SynCom at a time (Figure 6C,D). Interestingly, co-inoculation with wild-type WCS358 did not elevate either WCS358:pqqF or WCS358:cyoB abundance in the root compartment (Figure 6C). Thus, WCS358 wild-type did not *trans*-complement for the impaired root colonization of WCS358 mutants. On the contrary, in these mixed inoculation experiments (wild-type WCS358 plus WCS358 mutants), the abundance of wild-type WCS358 and to a greater extent that of WCS358:cyoB on roots decreased upon co-inoculation with WCS358:pqqF (Figure 6C), as was the case for most other SynCom members (Figure 5G,F). Furthermore, WCS358:cyoB was essentially outcompeted in the root and peat matrix when co-inoculated with wild-type WCS358 and WCS358:pqqF and the 15-member SynCom (Figure 6D). It is therefore unlikely that reduced gluconic acid and/ or 2-keto-gluconic acid levels explain the lower microbial load in the presence of the WCS358:pqqF mutant. Thus, although WCS358:pqqF and WCS358:cyoB mutants share defects in gluconic acid synthesis and suppression of plant immunity in mono-associations with *Arabidopsis* (Yu et al., 2019), the dominant impact on root-associated bacterial communities is restricted to WCS358:pqqF. This reveals previously unsuspected additional and distinct functions of *pqqF* compared to *cyoB* on roots. In summary, the MoBacTag allowed us to test for complementation or competition between strains encoding identical *16S rRNA* sequences during root microbiota establishment.

## Discussion

We have developed and validated DNA barcodes, which are co-amplified with natural endogenous bacterial *V5*–*V7 rRNA*, fungal *ITS* or plant *ITS* sequences by integrating respective primer binding sites into the DNA barcode. The MoBacTag enables direct tracking of multiple near-isogenic strains during conventional bacterial *16S rRNA* or fungal *ITS* community profiling. Since MoBacTag plasmids are based on modular cloning principles, each position within the multi-gene construct can be customized (Weber et al., 2011) by, for example, replacing *Tn7* elements with genomic regions for homologous recombination or inserting (*i*) additional community profiling primer binding sites into the DNA barcode, (*ii*) an expression cassette for complementation approaches, or (*iii*) available modular cloning-compatible elements such as intensity-optimized fluorescent markers (Geddes et al., 2018; Iverson et al., 2016).

Only a small number of genetic determinants in bacterial commensals influencing microbe–microbe and microbe–host interactions during plant microbiota establishment have thus far been identified (Getzke et al., 2023; Hemmerle et al., 2022; Liu et al., 2022). Bacterial DNA barcoding was applied to investigate antagonistic interactions of Pseudomonads leaf strains co-occurring on *Arabidopsis* (Shalev et al., 2022). However, the DNA barcode architecture used does not allow co-amplification with endogenous marker genes, meaning that community analysis is restricted to barcoded strains only. The MoBacTag barcodes were semi-randomly assembled by random ligation of synthesized oligonucleotides consisting of 454 barcodes to avoid homo- or dipolymer tracts (Figure 1). Unexpected PCR biases of individual 454-based barcodes required correction factors to determine the abundance of MoBacTag-labeled strains (Figure 2E,F). The possibility of a similar PCR bias being introduced by single 454 barcodes during standard library preparation for amplicon sequencing can be disregarded for compositional analysis of communities within a sample. However, amplicons of different samples are typically labeled with different 454 barcodes, which might introduce sample-specific PCR biases, complicating comparison of aggregated amplicon abundances between samples. We have used here a spike-in plasmid to calculate normalized abundances (Figure 2A–D), which eliminates potential 454 barcode- and thereby sample-specific PCR biases. Our spike-in plasmid architecture is similar to the architecture of other spike-in plasmids that require RT-qPCR quantification (Figure 1; Guo et al., 2020; Tkacz et al., 2018). Besides spike-in normalization, hamPCR-based approaches are also used to measure microbial load and community composition (Lundberg et al., 2021). MoBacTag and hamPCR approaches are complementary and could be combined to further increase the accuracy of host microbial load estimates.

MoBacTags are chromosomally integrated as a single copy, whereas *16S rRNA* gene copy numbers can vary from 1 to 15 between bacterial species (Louca et al., 2018; Vetrovsky & Baldrian, 2013). In this study, we compared only barcoded variants of a wild-type Rhizobiales Root13D or wild-type *P. capeferrum* WCS358 and its mutant derivatives for which potential *16S rRNA* copy number variation is irrelevant. In future experiments with multiple MoBacTag-labeled strains, representing different bacterial species in the community, the tag-specific correction factor must also be adjusted to the corresponding number of *16S rRNA* copies.

The spike-in normalization revealed a significantly reduced SynCom load in the presence of the WCS358:pqqF mutant, but not WCS358:cyoB, which can be explained by significantly decreased abundances of 7 of the 15 SynCom members, each representing different core taxonomic lineages of the *A. thaliana* root microbiota (Figure 5B,F,G; Bai et al., 2015; Wippel et al., 2021). WCS358:pqqF and WCS358:cyoB are both impaired in the acidification of the extracellular space, although the WCS358:cyoB mutant produces residual amounts of gluconic acid (Yu et al., 2019). As the WCS358:pqqF mutant, but not the WCS358:cyoB mutant, reduced microbial load on roots, which is not *trans*-complemented by gluconic-acid producing wild-type WCS358, the observed community shift is unlikely to be a direct consequence of impaired gluconic-acid-mediated host immunosuppression. We propose instead that the cofactor PQQ serves as a common good within the root-associated bacterial community, as has been shown for other cofactors such as cobamides in bacterial co-cultures composed of different species (Sokolovskaya et al., 2020). Extracellular PQQ levels are likely relevant in bacterial community contexts as PQQ import has been shown to be concentration-dependent: the cofactor is imported by diffusion at high concentrations, whereas active TonB-dependent import is required at low concentrations, at least for *E.coli* (Hantke & Friz, 2022). Thus, the WCS358:pqqF mutant could reduce the extracellular PQQ pool on roots due to lack of PQQ biosynthesis despite continuous PQQ consumption through WCS358:pqqF proliferation, resulting in PQQ deficiency in the bacterial community. This would also explain why WCS358:pqqF mutant-specific community phenotypes are not *trans*-complemented by wild-type *P. capeferrum* WCS358 *in planta*.

MoBacTag DNA barcodes generated *in vitro* were designed for tracking near-isogenic bacterial strains in community contexts across multiple generations. These strains cannot be distinguished by endogenous barcodes such as the *V5–V7 16S rRNA* region. The presence of the dominant-negative *ccdB* selection marker in the recipient vector (Figure 1) should enable use of MoBacTags for pool transformation approaches, each with randomly loaded DNA barcode tags (Figure 1; Kebschull & Zador, 2018). This opens up future opportunities for MoBacTag-based lineage tracking during experimental evolution of microbial communities (Venkataram et al., 2023), as potential compensatory growth responses of community members to changes in labeled lineages can be tracked in parallel. Although our proof-of-principle experiments were performed with SynComs, MoBacTag-labeled strains can be directly inoculated and tracked in environments harboring natural microbial communities, such as natural soils. Some MoBacTag-labeled strains may be barely competitive in highly complex microbial communities, such as in natural soil, making it unlikely that we can detect such a strain by routine endogenous *16S rRNA* amplicon sequencing. However, spike-normalized community profiling with barcode-specific primers allows strain detection in such cases as well as estimates of tag-to-plant *ITS* reads. Furthermore, replacement of primer binding sites for plant-specific marker gene amplification with binding sites for other eukaryotic hosts will increase the versatility of MoBacTag vectors for studies of any host-microbiota interaction.

## Material and Methods

### Bacterial Media and Growth Conditions

Level 1 vectors were transformed into *E. coli* DH5 and DB3.1, for *ccdB*-encoding plasmids (Supplemental table S1). Level 2 pBCC and pBC vectors were transformed into *E. coli* DB3.1λpir (House et al., 2004) and BW29427, also known as WM3064 (Dehio & Meyer, 1997), respectively (Supplemental table S1). pTNS3 (Choi et al., 2008) was also cloned into *E. coli* BW29427. *E. coli* were cultivated in LB (25 g.l^-1^ LB Broth, Sigma) medium at 37 °C, and commensal bacterial strains were cultivated in 0.5 TSB (15 g.l^-1^ Tryptic Soy Broth, Sigma) or TY (5 g.l^-1^ Tryptone, 3 g.l^-1^ Yeast extract, 10 mM CaCl_2_) medium at 25 °C. Media were supplemented with 15 g.l^-1^ Bacto Agar (Difco) for solidification. Antibiotics or diaminopimelic acid were added to the media, when necessary, at the following concentrations: streptomycin (Sm, 100 μg.ml^−1^), spectinomycin (Sp, 100 μg.ml^−1^) tetracycline (Tc, 10 μg.ml^−1^), gentamicin (Gm, 25 μg.ml^−1^), kanamycin (Kn, 50 μg.ml^−1^), and diaminopimelic acid (DAP, 50 μg.ml^−1^).

### Assembly of unique DNA barcodes by random ligation of oligonucleotides

Complementary oligonucleotides (Supplemental table S2) were mixed at an equimolar ratio (10 µM), incubated at 94 °C for 2 minutes and gradually cooled. Double-stranded oligonucleotides were then mixed in 1:1 ratios and prepared for blunt-end ligation using the End-It DNA End-Repair Kit according to the user manual (Biosearch Technologies). Ligation was performed overnight at 4 °C using T4 DNA ligase (New England Biolabs) followed by heat-inactivation for 10 min at 80 °C. The preassembled barcodes were finally selected for 200–300 bp fragments by BluePippin from Sage Science at the Max Planck Genome Centre, Cologne, Germany (https://mpgc.mpipz.mpg.de/home/). Size-selected arrays of ligated oligonucleotides were then cloned between primer binding sites of pBCC vectors (see next paragraph).

### Generation of MoBacTag plasmids using the Modular Cloning strategy (MoClo)

Expression cassettes, terminator sequences and mini*Tn7* elements were amplified using PrimerStar Max DNA Polymerase from Takara (templates indicated in supplemental table S3). PCR fragments were cloned into level 1 recipient plasmids (Supplemental table S3) from the MoClo ToolKit by a restriction/ligation reaction with *Bsa*I and T4 DNA ligase (New England Biolabs) as previously described (Weber et al., 2011; Werner et al., 2012). *Bsa*I, *Bpi*I and *Sma*I recognition sites were removed by altering a single nucleotide, without changing the encoded amino acid, in the recognition site using primers with restriction site-mutating sequences (Supplemental table 2, 3). The *ccdB* cassette was flanked with *Sma*I recognition sites and amplicon sequencing primer binding sites by consecutive PCRs with tailed primers (primers: Supplemental table S2) and then cloned into the level 1 recipient plasmid plCH47761. The antibiotic markers (pBCC030, pBCC031, pBCC032) were flanked with FRT sites by two consecutive PCRs using FRT site encoding tailed primers, whereas for pBCC033 the FRT sites were already present in the donor plasmid (Supplemental table S3). First, a GFP-encoding plasmid was assembled (pBCC029) by amplifying GFP from pUC18-mini-Tn7T-Tp-gfpmut3 and pTAC from pLM449 with tailed primers integrating *Bsa*I restriction sites and cloned into the plCH47751 Level 1 receptor as previously described. Finally, the pTAC promoter and coding sequence of the fluorescent markers were amplified from pLM426 derivatives (Supplemental table S3) and combined with the pBCC0029 recipient plasmid including a terminator sequence by In-Fusion HD from Takara. All plasmids for this study were purified using the NucleoSpin Plasmid kit (Macherey-Nagel). Level 1 domesticated sequences were then assembled into a pSEVA211-based (Silva-Rocha et al., 2013) level 2 recipient vector by a *Bpi*I restriction/ligation reaction resulting in BarCode Construction (pBCC) multi-gene constructs (Supplemental table S3). To this end, a tagRFP expression cassette was amplified by PCR integrating *Bpi*I recognition sites and cloned into the pSEVA211 by a restriction/ligation reaction with *Bsa*I, *Eco*RI, *Hind*III and T4 DNA ligase (New England Biolabs). Pre-assembled barcodes were inserted by *Sma*I restriction/ligation as follow: 45 cycles of 5 min at 16 °C; 5 min at 30 °C followed by 5 min 95 °C. Then, fresh *Sma*I was added and incubated at 30 °C for 30 min resulting in final MoBacTag vectors. Barcode tag sequences were identified for individual *E. coli* BW29427 colonies from pooled transformation with MoBacTag vectors by colony PCR with primers MobacTag barcode F/R (Supplemental table S5), followed by Sanger sequencing (Eurofins Scientific). Sequencing results were analyzed in CLC Main Workbench (QIAGEN).

### Labeling of bacterial strains with MoBacTags using mini *Tn7* system

MoBacTag vectors were transferred into commensal bacterial strains (Supplemental table S1) by triparental mating. Saturated liquid cultures of the recipient commensal strain, *E. coli* strain BW29427/pTNS3 and BW29427/pBC were mixed in a 1:2:2 ratio and incubated for 24–48h at 25 °C. Afterwards, transformants were selected on 0.5 TSB or TY medium containing 10% sucrose and the corresponding antibiotics. Bacterial DNA was extracted by re-suspending a bacterial colony in 25 µl buffer I (25 mM NaOH, 0.2 mM EDTA, pH 12) followed by the incubation at 95 °C for 30 min and addition of 25 µl buffer II (40 mM Tris-HCl). The genomic insertion of the MoBacTag was validated by PCR as described in Choi and Schweizer (2006).

### Bacterial microbiota reconstitution experiments

Saturated bacterial liquid cultures were pelleted by centrifugation at 8,000 g for 5 min, followed by two washes with 10 mM MgSO_4_. Equivalent amounts of each strain were combined to yield the desired SynComs with an optical density (OD_600_) of 2. Aliquots of individual strains and the SynComs were taken and stored at −80 °C. The inoculum solution was prepared with MS/2 (2.22 g.l^−1^ Murashige+Skoog basal salts, Sigma; 0.5 g.l^−1^ MES anhydrous, BioChemica; adjusted to pH 5.7 with KOH) and the SynCom at a final OD_600_ of 0.02. We used the Gnotopot system (Kremer et al., 2021) to grow *A. thaliana* Col-0 plants with the bacterial SynComs (Supplemental table S6). Each pot was inoculated with the bacterial SynCom by decanting 10 ml of the inoculum solution. With the help of a syringe, the excess of liquid was removed from the box. *A. thaliana* seeds were surface-sterilized by incubation in 70% ethanol twice for 5 min, followed by a brief wash with 100% ethanol. Seeds were then washed three times with sterile water and cold-stratified for two days. Six sterilized seeds were placed on the matrix of each pot (Jiffy-7 pellets, manufactured by Jiffy Products, Norway, https://www.jiffygroup.com/) and incubated under short-day conditions for five weeks (10 h light, 21 °C; 14 h dark, 19 °C). Roots were harvested by thoroughly cleaning from attached soil using sterile water and forceps. Root and peat matrix samples were collected in Lysing Matrix E tubes (FastDNA Spin Kit for Soil, MP Biomedicals) and frozen in liquid nitrogen. Samples were stored at −80 °C until DNA isolation, which was performed using the FastDNA Spin Kit for Soil according to the user manual (MP Biomedicals).

### Visualization of fluorescently labeled commensal bacteria

MoBacTag-labeled bacterial strains were harvested from 0.5 TSB plates and re-suspended in 10 mM MgSO_4_. The root colonization assay was performed as follows: surface-sterilized *A. thaliana* Col-0 seeds were sown on agar plates (1% Bacto agar BD Biosciences) prepared with MS/2 (as described for the Gnoptopot system) and supplemented with MoBacTag-labeled *Rhodanobacter* Root179 or Rhizobiales Root13D at a final concentration of OD_600_ = 0.0005 (Ma et al., 2022). Washed bacteria (see bacterial microbiota reconstitution experiment) were mixed into the medium prior to solidification. After 14 days of growth under short-day conditions (10 h light, 21 °C; 14 h dark, 19 °C), bacteria were visualized on roots. Confocal laser scanning microscopy was performed on a Zeiss LSM880 inverted confocal scanning microscope. Pictures were taken with a LD C-Apochromat 40x/1.1 water immersion objective. To image root colonization, Z-stacks were generated, and maximum intensity projections were compiled. The following excitation and detection windows were used: GFP 488 nm, 493-598 nm; AmCyan1 458 nm, 472-528 nm; tagRFP 561 nm, 582-754 nm; far-red-FP 561 nm, 591-759 nm.

### Bacterial community profiling by amplicon sequencing

Library preparation for Illumina MiSeq sequencing was performed as described previously (Duran et al., 2018), except for adding 0.001 ng of pBCC023 or pBCC084 plasmid DNA per reaction to the master mix of PCR1, as spike. The final ratio of sample (6 ng) to spike (0.001 ng) DNA per PCR1 reaction is 6,000. The list of oligonucleotides used for amplicon sequencing of *16S rRNA*, fungal *ITS* and plant *ITS* sequences are listed in supplemental table S5. In all experiments, multiplexing of samples was performed by single or double-indexing (only forward barcoded oligonucleotides for single indexing or forward and reverse barcoded oligonucleotides for double-indexing). The indexed amplicon libraries were pooled, purified using Ampure (Beckman-Coulter) and sequenced on the Illumina MiSeq platform. To validate MoBacTag as a spike-in plasmid using the bacterial *16S*-, fungal *ITS*-, plant *ITS*- and barcode-specific primers, 0.15 ng of pBCC069 plasmid were mixed with 5 ng of extracted DNA from root and peat matrix samples, prepared by Getzke et al. (2023). The standard curve was prepared with pBCC084 plasmid in tenfold dilution series from 1.5 to 0.00015 ng per PCR reaction.

### Processing of gene amplicon data

Amplicon sequencing data from SynCom experiments were de-multiplexed according to their barcode sequence and quality-filtered using the QIIME pipeline (Caporaso et al., 2010). Paired-end reads were merged using the flash2 software (Magoc & Salzberg, 2011). Quality-filtered merged paired-end reads were then aligned to reference amplicon sequences using Rbec (v1.0.0) (Zhang et al., 2021). For the plant *ITS* sequencing data, only single-end reads were processed, because sequence length of pITS (742 bp) can’t be merged from Illumina 2 X 300 bp sequencing run. The reference sequences were extracted from whole-genome assemblies of every strain included in the SynCom, from the MoBacTag barcodes and from whole-genome assembly of *A. thaliana* Col-0 (TAIR9 assembly, www.arabidopsis.org). We checked that the fraction of unmapped reads did not significantly differ between compartments and experiments. Count tables were generated and employed for downstream analyses of diversity in R using the R package vegan (Dixon, 2003). Amplicon data were visualized using the ggplot2 R package (Villanueva & Chen, 2019).

### Normalized quantification of amplicon sequencing

Amplicon reads assigned to the spike were used to normalize plant ITS and bacterial 16S rRNA read counts similarly to Tkacz et al. (2018). Identical amounts of spike-in plasmid DNA included for plant ITS and 16S rRNA library preparation were used for cross-normalization using the following equations:

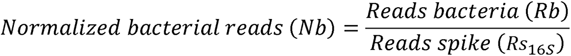

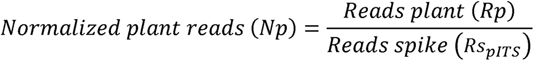

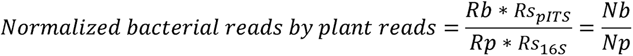

## Statistics and reproducibility

All experiments were performed in three full factorial (biological and technical) replication. Bacterial abundances were compared using an ANOVA test, followed by a Tukey’s post hoc (α = 0.05). Statistical tests on beta-diversity analyses were performed using a PERMANOVA test with 999 random permutations. Whenever boxplots were used in figures, data were represented as median values (horizontal line), Q1 − 1.5× interquartile range (boxes) and Q3 + 1.5× interquartile range (whiskers).

## Data availability

Raw amplicon reads and genome assemblies have been deposited in the European Nucleotide Archive under the accession number PRJEB61076. The plasmids generated for the MoBacTag kit will be deposited to Addgene.

## Acknowledgments

We thank D. Becker for technical support. We acknowledge P. Poole for providing the pLM449 plasmid containing the tac promoter and F. Getzke for providing root and peat matrix samples containing fungal DNA. We thank N. Donnelly, J. Stuttmann and R. Berendsen for reading and editing the manuscript. Funding was provided by the Max Planck Society and the German Research Foundation (DFG) under the German Excellence Strategy, EXC number 2048/1 project 390686111 for R.G.-O. and P.S.-L., and SPP 2125 DECRyPT for K.-W.M. and P.S.-L. J.T. was supported by the Alexander von Humboldt Foundation.

## Author contributions

P.S.-L., J.O., K.-W.M., R.G.-O. and B.H. conceptualized the methodology. J.O. and J.T. designed the experiments. R.T.N. cloned coding sequences for fluorophores. J.O. and J.T. generated MoBacTag plasmids. J.O. assembled barcode DNA tags and tested their diversity. J.T. tested spike-in normalization and amplification biases by multiple MoBacTag-tagged strains. J.O. performed microscopy and the proof-of-principle experiment. J.O., J.T. and P.Z. analyzed the data. J.O. and J.T. produced the figures. J.O. and P.S.-L. wrote the manuscript with contributions from all co-authors.

## Supplemental figure legends

**Supplemental figure 1:**
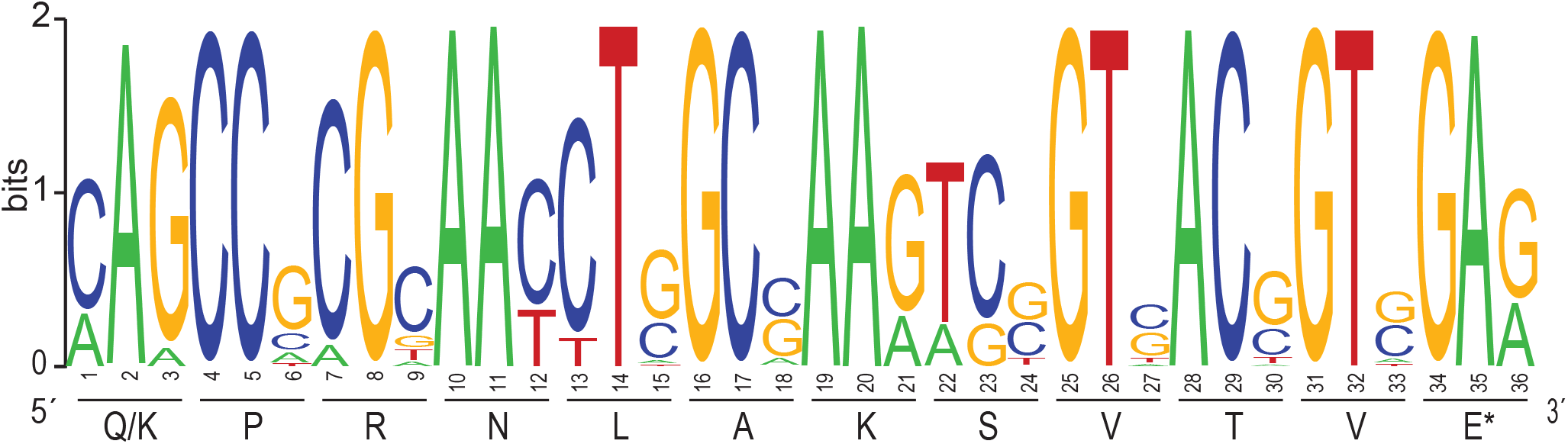
Conservation of the *Tn7* binding site in the *At*-SPHERE culture collection. *Tn7* transposase binding site within the last coding 36 nts of the *glucosamine-6-phosphate synthetase* (*glmS*) gene, displayed as a DNA sequence logo for all *At*-SPHERE genome drafts. The letter size indicates the nucleotide abundance at each position. The encoded amino acids are depicted below.

**Supplemental figure 2:**
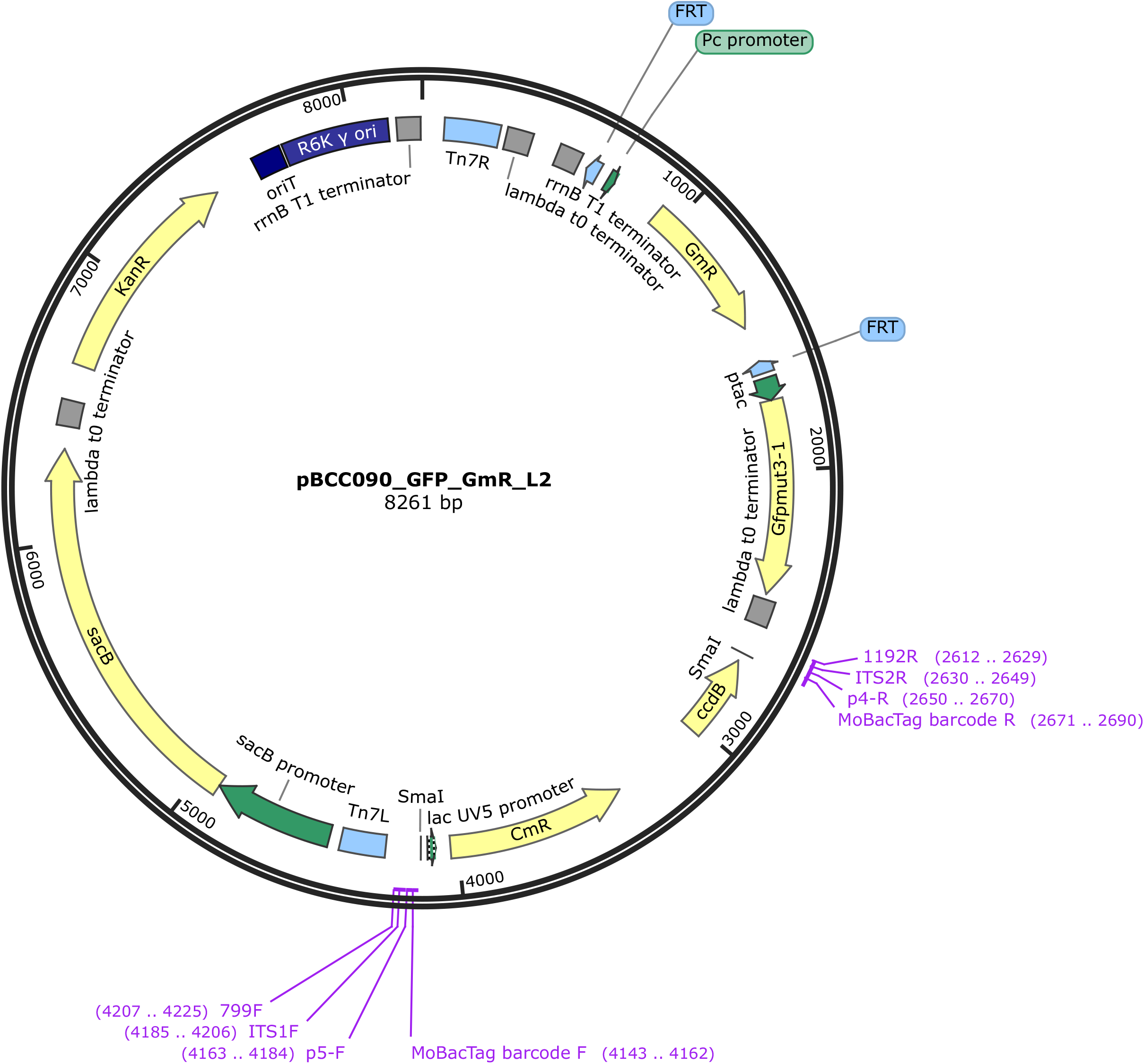
Plasmid map of an *N-acetyltransferase aac* and *GFP*-encoding pBCC recipient plasmid. Plasmid map, created with SnapGene, indicating coding sequences (yellow), promoter sequences (green), terminator sequences (grey), origin of transfer and replication (dark blue), protein binding sites (light blue), primer binding sites (purple) and *Sma*I restriction sites.

**Supplemental figure 3:**
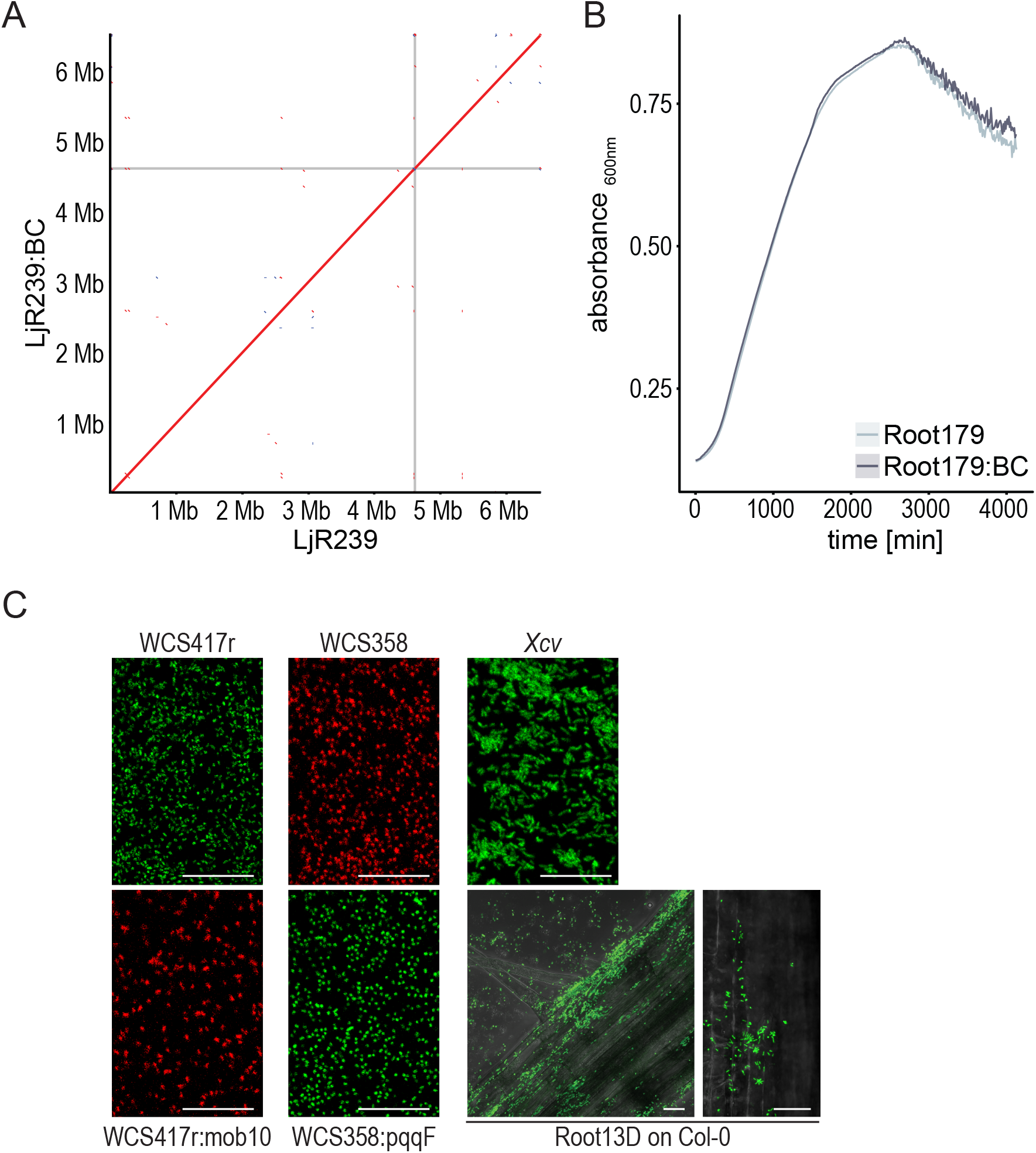
Chromosome integrity and expression of fluorescent tags in taxonomically distinct bacteria. (A) Alignments of PacBio genome assemblies from the unlabeled and MoBacTag-labeled Rhizobiales LjR239 isolated from *Lotus japonicus* roots. MoBacTag insertion site is indicated by grey lines. (B) Bacterial growth of MoBacTag-labeled or unlabeled *Rhodanobacter* Root179 in liquid rich (0.5 TSB) medium in monocultures indicated by absorbance (λ = 600nm). (C) Expression of fluorescent proteins from chromosomally-integrated MoBacTags. Expression of MoBacTag-encoded fluorescent protein was detected using live confocal laser scanning microscopy in liquid medium and on *A. thaliana* roots. Xcv: *Xanthomonas campestris* pv. *vesicatoria*. Scale: 20 µm.

**Supplemental figure 4:**
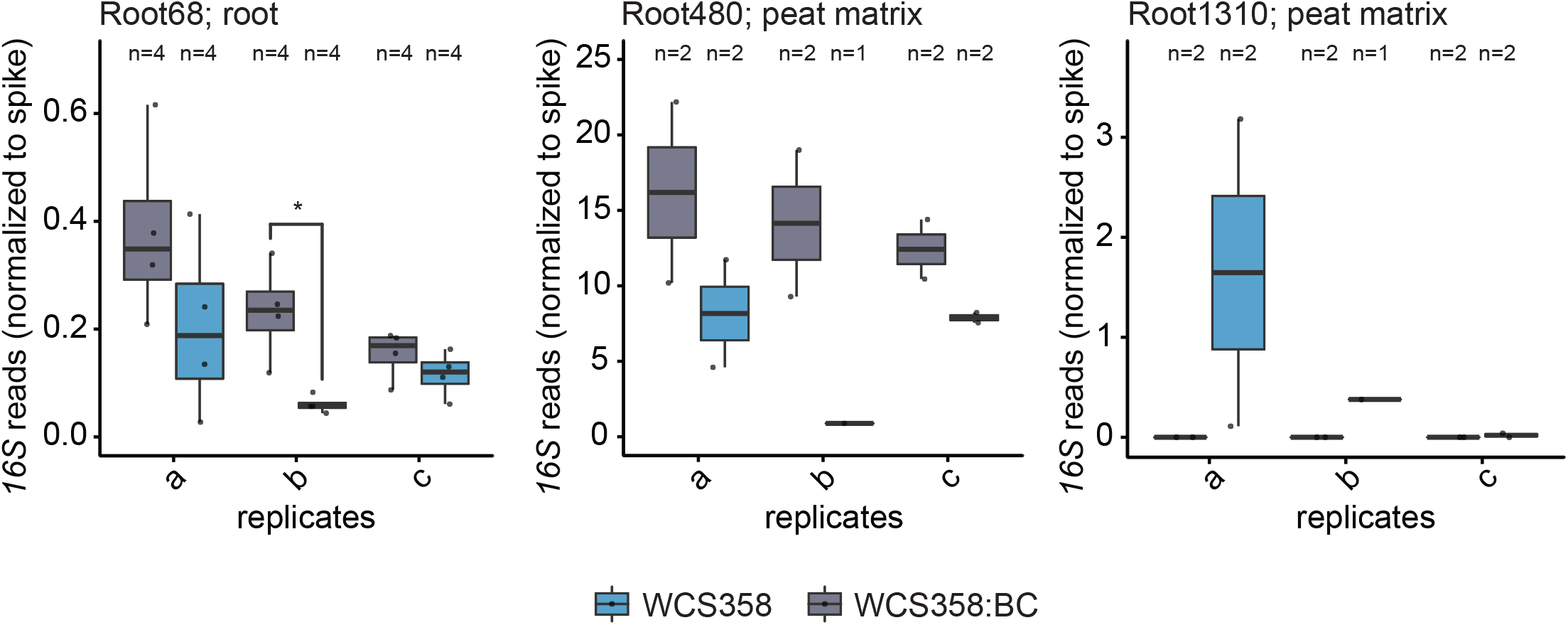
The influence of a MoBacTag on selected SynCom members in independent experiments. The normalized abundance of strains that had significantly different abundances after co-inoculation with the labeled (WCS358:BC) or unlabeled (WCS358) strain when all replicates are analyzed collectively (Figure 4 D,E) for each independent experiment (a, b, c).

## Supplemental Material and Methods

### *AttTn7* box conservation across At-SPHERE culture collection

The *glmS* sequences were extracted from the whole-genome assemblies of every strain included in the At-SHPERE culture collection (Bai et al., 2015). The last 12 amino acids or the last 36 DNA bases from the extracted *glmS* sequences were aligned and visualized with the software WebLogo (https://weblogo.berkeley.edu/logo.cgi)

### Bacterial genome assembly

The Max Planck Genome Centre, Cologne, Germany (https://mpgc.mpipz.mpg.de/home/) performed the DNA isolation of wild-type and MoBacTag-labeled strains and also the sequencing on the Pacific Biosciences Sequel IIe platform. Reads were assembled *de novo* using the Hifiasm software (https://github.com/chhylp123/hifiasm). To compare wild-type and MoBacTag-labeled strains, we used Mauve software for reordering the contigs (https://darlinglab.org/mauve). Genomes were compared by generating a dotplot on genome scale using the Genome Pair Rapid Dotter (gepard) software (https://doi.org/10.1093/bioinformatics/btm039).

### Bacterial growth in liquid mono-culture

MobacTag-labeled and unlabeled *Rhodanobacter* R179 strains were grown in six replicates each in 0.5 TSB as individual cultures in a 96-well bacterial culture plate (Greiner-CELLSTAR 96-well plate, transparent, flat bottom; Sigma-Aldrich) at 25 °C. Absorbance was measured every 10 minutes, 10 seconds after 20 seconds of shaking (290 rpm) using a microplate reader (Infinite M200 PRO, Tecan) at 600 nm. The mean value of four measurements per well was used for the analyses.

